# Prediction of prostate cancer by deep learning with multilayer artificial neural network

**DOI:** 10.1101/291609

**Authors:** Takumi Takeuchi, Mami Hattori-Kato, Yumiko Okuno, Satoshi Iwai, Koji Mikami

**Affiliations:** Department of Urology, Japan Organization of Occupational Health and Safety, Kanto Rosai Hospital, 1-1 Kizukisumiyoshi-cho, Nakahara-ku, Kawasaki 211-8510, Japan; Department of Medical Informatics and Economics, Graduate School of Medicine, The University of Tokyo, Japan, 7-3-1 Hongo, Bunkyo-ku, Tokyo 113-8654, Japan

**Keywords:** prostate, biopsy, artificial intelligence, neural network

## Abstract

**Objectives:** To predict the rate of prostate cancer detection on prostate biopsy more accurately, the performance of deep learning utilizing a multilayer artificial neural network was investigated.

**Materials and methods:** A total of 334 patients who underwent multiparametric magnetic resonance imaging before ultrasonography-guided transrectal 12-core prostate biopsy were enrolled in the analysis. Twenty-two non-selected variables as well as selected ones by least absolute shrinkage and selection operator (Lasso) regression analysis and by stepwise logistic regression analysis were input into the constructed multilayer artificial neural network (ANN) programs. 232 patients were used as training cases of ANN programs, and the remaining 102 patients were for the test to output the probability of prostate cancer existence, accuracy of prostate cancer prediction, and area under the receiver operating characteristic (ROC) curve with the learned model.

**Results:** With any prostate cancer objective variable, Lasso and stepwise regression analyses selected 12 and 9 explanatory variables from 22, respectively. In common between them, age at biopsy, findings on digital rectal examination, findings in the peripheral zone on MRI diffusion-weighted imaging, and body mass index were positively influential variables, while numbers of previous prostatic biopsy and prostate volume were negatively influential. Using trained ANNs with multiple hidden layers, the accuracy of predicting any prostate cancer in test samples was about 5-10% higher compared with that with logistic regression analysis (LR). The AUCs with multilayer ANN were significantly larger on inputting variables that were selected by the stepwise logistic regression compared with the AUC with LR. The ANN had a higher net-benefit than LR between prostate cancer probability cut-off values of 0.38 and 0.6.

**Conclusion:** ANN accurately predicted prostate cancer without biopsy marginally better than LR. However, for clinical application, ANN performance may still need improvement.

## Introduction

Prostate cancer was the third most common cancer diagnosed in men and the sixth most common cause of cancer death in Japan in 2017 (1). Prostatic biopsy is indicated when a patient is suspected of having prostate cancer, mainly based on elevated serum prostate-specific antigen (PSA) and/or digital rectal examination (DRE). Nevertheless, the rate of detecting prostatic cancer upon biopsy was 32% based on the Surveillance, Epidemiology, and End Results (SEER) data (2). Recently, MRI-targeted biopsy enhanced the rate of detecting any prostate cancer to 51-66% (3–6). If the existence of prostate cancer can be predicted more accurately before biopsy, unnecessary and invasive prostatic biopsy can be avoided.

Recent advances in artificial intelligence are now being applied to various fields in society and science. A neural network is worth mathematically modeling the brain by mimicking its pattern recognition capabilities. It has strength in non-linear classification including complex regression problems. Here, we have predicted the rate of prostate cancer detection on prostate biopsy by deep learning utilizing a multilayer artificial neural network (ANN) as an application to the urologic field.

## Materials and Methods

A total of 334 patients who underwent 3-Tesla multiparametric magnetic resonance imaging (mpMRI) before ultrasonography-guided MRI-targeted transrectal 12-core prostate biopsy between 2013 and 2017 were enrolled in the analysis. MRI was viewed before biopsy and the MRI-identified lesion was cognitively targeted. Upon biopsies, the sample sites from the prostatic peripheral zone were bilateral from the apex, mid-gland, to base. Two additional cores from the far posterior, lateral peripheral zone and a core from the transition zone of each lobe were also biopsied. Exceptionally, seven patients with PSA values between 32-725 ng/ml, positive DRE, and positive MRI findings were administered sextant biopsy and prostate cancer was detected in all. Patient and clinical characteristics are described in Table 1.

**Table 1:** Clinical characteristics

|  |  |  |  |  |  |  |
| --- | --- | --- | --- | --- | --- | --- |
|  | Median (Range) |  | Median (Range) |  |  |  |
| Age: years | 70 (44-88) | TG: mg/dL | 126 (35-497) |  |  |  |
| BH: cm | 167 (152-180) | BS: mg/dL | 107 (72-278) |  |  |  |
| BW: kg | 64 (39-111) | eGFR: mL/min | 68 (3.9-110) |  |  |  |
| BMI | 24 (14-37) | GPT: unit/L | 20 (4-78) |  |  |  |
| WBC: /μL | 5,800 (3,000-13,900) | CRP: mg/dL | 0.06 (0.003-4.45) | | | |
| Hb: g/dL | 14.4 (6.9-19.9) | PSA: ng/mL | 8.8 (3.5-725) |  |  |  |
| Alb: mg/dL | 4.4 (2.2-5.0) | PV: mL | 30 (10-274) |  |  |  |
| Tchol: mg/dL | 197 (85-308) | PSAD | 0.3 (0.04-47) |  |  |  |
|  | N (%) |  |  |  |  |  |
| DRE | 59 (18) |  |  |  |  |  |
| MRI_PZ_T2-positive | 157 (47) |  |  |  |  |  |
| MRI_PZ_DWI-positive | 154 (46) |  |  |  |  |  |
| MRI_TZ_T2-positive | 131 (39) |  |  |  |  |  |
| MRI_TZ_DWI-positive | 130 (39) |  |  |  |  |  |
| Previous biopsIES | 0 | 1 | 2 | 3 | 4 | 5 |
| N (%) | 287 (86) | 31 (9) | 9 (3) | 6 (2) | 0 (0) | 1 (0) |
| GS of biopsy-detected Pca | 6 | 7 | 8 | 9 | 10 |  |
| N (%) | 51 (15) | 37 (11) | 53 (16) | 24 (7) | 4 (1) |  |

Age: age at prostate biopsy, previous biopsies: number of prostate biopsies performed previously, BH: body height, BW: body weight, BMI: body mass index, WBC: white blood cell counts in blood, Hb: hemoglobin concentration, Alb: serum albumin, Tchol: serum total cholesterol, TG: serum triglycerides, BS: blood sugar, eGFR: estimated glomerular filtration rate, GPT: serum glutamic pyruvic transaminase, CRP: serum C-reactive protein, PSA: PSA before biopsy, PV: prostate volume calculated by trans-abdominal ultrasonography, PSAD: PSA density, DRE: digital rectal examination, PZ: prostatic peripheral zone, TZ: transition zone, T2: T2-weighted imaging, DWI: diffusion-weighted imaging, DRE and MRI findings were recorded as 1 if prostate cancer was suspected, otherwise 0, Pca: prostate cancer, GS: Gleason score, N: Patient number

The library to construct deep leaning programs was tensorflow (Google) and the programming language was python. Twenty-two variables (Table 2) of patients who underwent prostate biopsy were initially enrolled with continuous variables standardized ((value-average)/standard_deviation). Appropriate variables were selected by least absolute shrinkage and selection operator (Lasso) regression analysis (7) and by stepwise logistic regression analysis using data from all 334 samples with the glmnet 2.0-10 and the glm packages for R version 3.2.4 (Table 2). Then, 22 non-selected variables as well as automatically selected ones were input into the constructed multilayer artificial neural network (ANN) programs and were also analyzed with the conventional logistic regression analysis (LR). In brief, 232 patients with digits 0-6 in the one’s place of patient identification (ID) numbers were used as training cases in ANN programs, and the remaining 102 patients with 7-9 there were for the test to output the probability of prostate cancer detection, accuracy of prostate cancer prediction, and area under the receiver operating characteristic (ROC) curve with the learned model. When the output probability of prostate cancer detection in a patient was more than 0.5, it was regarded as prostate cancer and vice versa, as a discrete classifier. The parameter of ROC curves was the probability of prostate cancer detection to be evaluated as a probabilistic classifier.

**Table 2:** Variable selection by the Lasso and stepwise regression analyses.

|  | Lasso coef | Stepwise coef |  | Lasso coef | Stepwise coef |
| --- | --- | --- | --- | --- | --- |
| (Intercept) | 0.338 | -0.050 | eGFR | . | . |
| Age | 0.047 | 0.461 | GPT | . | . |
| Previous biopsies | -0.070 | -0.740 | CRP | . | . |
| BH | . | 0.741 | PSA | 0.002 | . |
| BW | . | -1.189 | PV | -0.040 | -0.245 |
| BMI | 0.006 | 1.319 | PSAD | . | 3.835 |
| WBC | . | . | DRE | 0.269 | 2.139 |
| Hb | . | . | MRI_PZ_T2 | . | . |
| Alb | . | . | MRI_PZ_DWI | 0.258 | 1.031 |
| Tchol | -0.009 | . | MRI_TZ_T2 | 0.027 | . |
| TG | -0.001 | . | MRI_TZ_DWI | 0.016 | . |
| BS | -0.008 | . |  |  |  |

Lasso coef and Stepwise coef were coefficients of remaining variables by Lasso and stepwise regression analysis, respectively, dot: absent variables

Two to five hidden layers of ANN were composed of 5 neurons for each layer, the activation function of hidden layers was the ReLu function, loss function was cross entropy error function, back propagation algorithm was gradient descent, and output function was softmax function. Validation of the training of ANNs was performed with the 5-fold cross-validation method (Figure 1). The learning rate of each step was 0.0001 and the L2 regularization penalty of scale 0.005 was added to cross entropy to avoid over-fitting; those values had been adopted in a preliminary evaluation of ANNs.

**Fig. 1:**
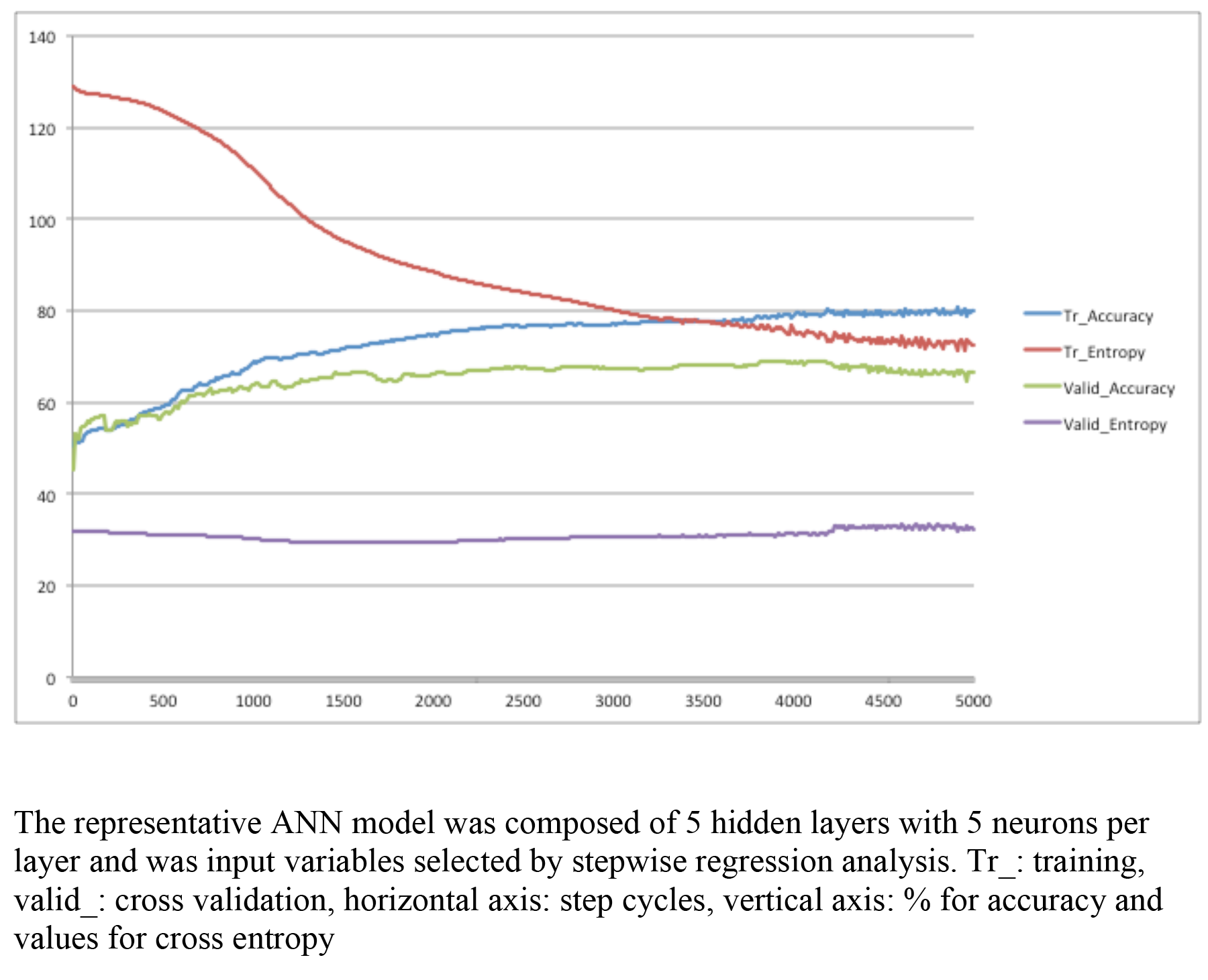
Accuracy and cross entropy in training and cross-validation of the ANN model.

LR with 22 non-selected variables and variables selected by Lasso and stepwise regression was also performed with R 3.2.4 and the glm package, inputting variables of the training samples described above (Table 3). Using the obtained coefficients, the probability of prostate cancer detection, accuracy of prostate cancer prediction, and area under the ROC curve (AUC) were output from test samples. When the probability of prostate cancer detection in a patient was more than 0.5, it was regarded as prostate cancer and vice versa, as described above.

**Table 3:** Logistic regression analysis of training samples with non-selected 22 variables and variables selected by the Lasso and stepwise regression.

| Coefficients: | All | Lasso | Stepwise |
| --- | --- | --- | --- |
| (Intercept) | 0.51±0.65 | 0.23±0.56 | 0.84±0.61 |
| Age | 0.43±0.28 | 0.17±0.20 | 0.29±0.21 |
| Previous biopsies | -0.58±0.34 | -0.60±0.34 | -0.55±0.32 |
| BH | -0.10±0.48 |  | -0.07±0.41 |
| BW | 0.68±0.87 |  | 0.16±0.74 |
| BMI | 0.19±0.76 | 0.61±0.21** | 0.46±0.65 |
| WBC | 0.23±0.22 |  |  |
| Hb | -0.42±0.29 |  |  |
| Alb | 0.41±0.25 |  |  |
| Tchol | -0.03±0.20 | -0.03±0.19 |  |
| TG | -0.09±0.21 | -0.14±0.2 |  |
| BS | -0.21±0.20 | -0.11±0.17 |  |
| eGFR | 0.22±0.20 |  |  |
| GPT | 0.12±0.23 |  |  |
| CRP | -0.48±0.31 |  |  |
| PSA | 3.87±3.78 | 6.46±2.19** |  |
| PV | -0.17±0.24 | -0.26±0.15 | 0.07±0.15 |
| PSAD | 5.67±5.65 |  | 10.17±3.25** |
| DRE | 3.87±1.15*** | 3.49±1.08** | 3.36±1.07** |
| MRI_PZ_T2 | -0.38±0.92 |  |  |
| MRI_PZ_DWI | 2.08±0.92* | 1.66±0.38*** | 1.27±0.35*** |
| MRI_TZ_T2 | -0.86±0.83 | -0.41±0.76 |  |
| MRI_TZ_DWI | 1.54±0.85 | 1.03±0.76 |  |

Values: estimate ± standard error, \*: p<0.05, \*\*: p<0.01, \*\*\*: p<0.001, All: 22 non-selected variables, Lasso: variables selected by Lasso regression analysis, Stepwise: variables selected by stepwise regression analysis

ROC curves were constructed and depicted using R 3.2.4 with the ROCR package for prostate cancer detection upon biopsy to assess the usefulness of the algorithm-output probability of prostate cancer detection as a parameter, and AUCs were statistically compared by R 3.2.4 with the pROC package. Net-benefit curves were depicted by R 3.2.4 with the rmda package.

Ethics: This study was conducted in accordance with the Helsinki declaration after approval by the Ethics Committee of Kanto Rosai Hospital.

## Results

### Variable selection with Lasso and stepwise regression analyses

With any prostate cancer objective variable, Lasso and stepwise regression analyses selected 12 and 9 explanatory variables from 22, respectively, as shown in Table 2. In common between them, age at biopsy, findings on DRE, findings in the peripheral zone on MRI diffusion-weighted imaging (MRI_PZ_DWI), and body mass index (BMI) were positively influential variables, while numbers of previous prostatic biopsy and prostate volume were negatively influential.

### Prediction of prostate cancer with multilayer ANN

Using trained ANNs with multiple hidden layers, the accuracy of predicting any prostate cancer in test samples was about 5-10% higher compared with that with LR, as shown in Table 4 (p-value not below 0.05 by Pearson chi-square test compared with corresponding LR, Odds Ratio 1.49). Variable selection with Lasso and stepwise regression analyses enhanced the accuracy of prostate cancer prediction and AUCs. Five thousand learning steps reduced the accuracy compared with 2,000 steps, possibly due to over-fitting. The AUCs with multilayer ANN were significantly larger on inputting variables that were selected by the stepwise logistic regression compared with the AUC with LR. The number of hidden layers slightly enhanced the AUCs.

**Table 4:** Performance of multilayer artificial neural network and logistic regression analysis.

| Any prostate cancer prediction in test samples |  |  |  |  |  |  |
| --- | --- | --- | --- | --- | --- | --- |
|  | Test_Set_Accuracy (%) |  |  | Area under ROC curve |  |  |
| Analytical method | Step1000 | Step2000 | Step5000 | Step1000 | Step2000 | Step5000 |
| 22_variable_Logistic_regression | 59.8 | 59.8 | 59.8 | 0.67 | 0.67 | 0.67 |
| 22_variable_ANN_2_hidden_layers | 66.7 | 64.7 | 57.8 | 0.72 | 0.70 | 0.64 |
| 22_variable_ANN_3_hidden_layers | 66.7 | 64.7 | 55.9 | 0.71 | 0.66 | 0.62 |
| 22_variable_ANN_4_hidden_layers | 65.7 | 60.8 | 57.8 | 0.70 | 0.66 | 0.61 |
| 22_variable_ANN_5_hidden_layers | 66.7 | 63.7 | 59.8 | 0.71 | 0.67 | 0.62 |
| Lasso_Logistic_regression | 61.8 | 61.8 | 61.8 | 0.68 | 0.68 | 0.68 |
| Lasso_ANN_2_hidden_layers | 68.6 | 70.6 | 69.6 | 0.74 | 0.70 | 0.69 |
| Lasso_ANN_3_hidden_layers | 68.6 | 71.6 | 64.7 | 0.75 | 0.72 | 0.69 |
| Lasso_ANN_4_hidden_layers | 67.7 | 71.6 | 68.6 | 0.73 | 0.72 | 0.70 |
| Lasso_ANN_5_hidden_layers | 66.7 | 71.6 | 67.7 | 0.73 | 0.71 | 0.72 |
| Stepwise_Logistic_regression | 61.8 | 61.8 | 61.8 | 0.68 | 0.68 | 0.68 |
| Stepwise_ANN_2_hidden_layers | 65.7 | 65.7 | 61.8 | 0.74 | 0.72 | 0.71 |
| Stepwise_ANN_3_hidden_layers | 64.7 | 68.6 | 62.8 | 0.73 | 0.75* | 0.67 |
| Stepwise_ANN_4_hidden_layers | 62.8 | 68.6 | 66.7 | 0.73 | 0.75* | 0.72 |
| Stepwise_ANN_5_hidden_layers | 65.7 | 70.6 | 67.7 | 0.74 | 0.76* | 0.74 |

Test\_Set\_Accuracy: prediction accuracy of test samples by ANN and LR when threshold set at 0.5, Step: step cycles of ANN, ignore for LR, 22\_variable: 22 non-selected variables, Lasso: variables selected by Lasso regression analysis, Stepwise: variables selected by stepwise regression analysis, \*: $p < 0.05$ by pROC compared with corresponding LR

The multilayer ANN that showed the largest AUC, i.e., ANN with 5 hidden layers using variables selected by stepwise regression analysis after 2,000 steps (Stepwise_ANN_5_hidden_layers in Table 4, its ROC curve in Figure 2), was precisely compared with the corresponding LR, as shown in Table 5. With a prostate cancer probability threshold of 0.3 for example, the ANN prevented 48% of patients without actual prostate cancer from having to undergo needless prostate biopsy, while LR prevented 44%. The ANN missed 16% of any prostate cancer and 6% of prostate cancer with GS≥7, while LR missed 18 and 9%, respectively. Negative predictive values (NPV) for any prostate cancer were 76 and 72% with ANN and LR, respectively. NPVs for prostate cancer with GS≥7 were 94 and 91%, respectively. As shown in Figure 3, net-benefit curves for any prostate cancer detection indicated that the ANN had a higher net-benefit than LR between prostate cancer probability cut-off values of 0.38 and 0.6.

**Fig. 2:**
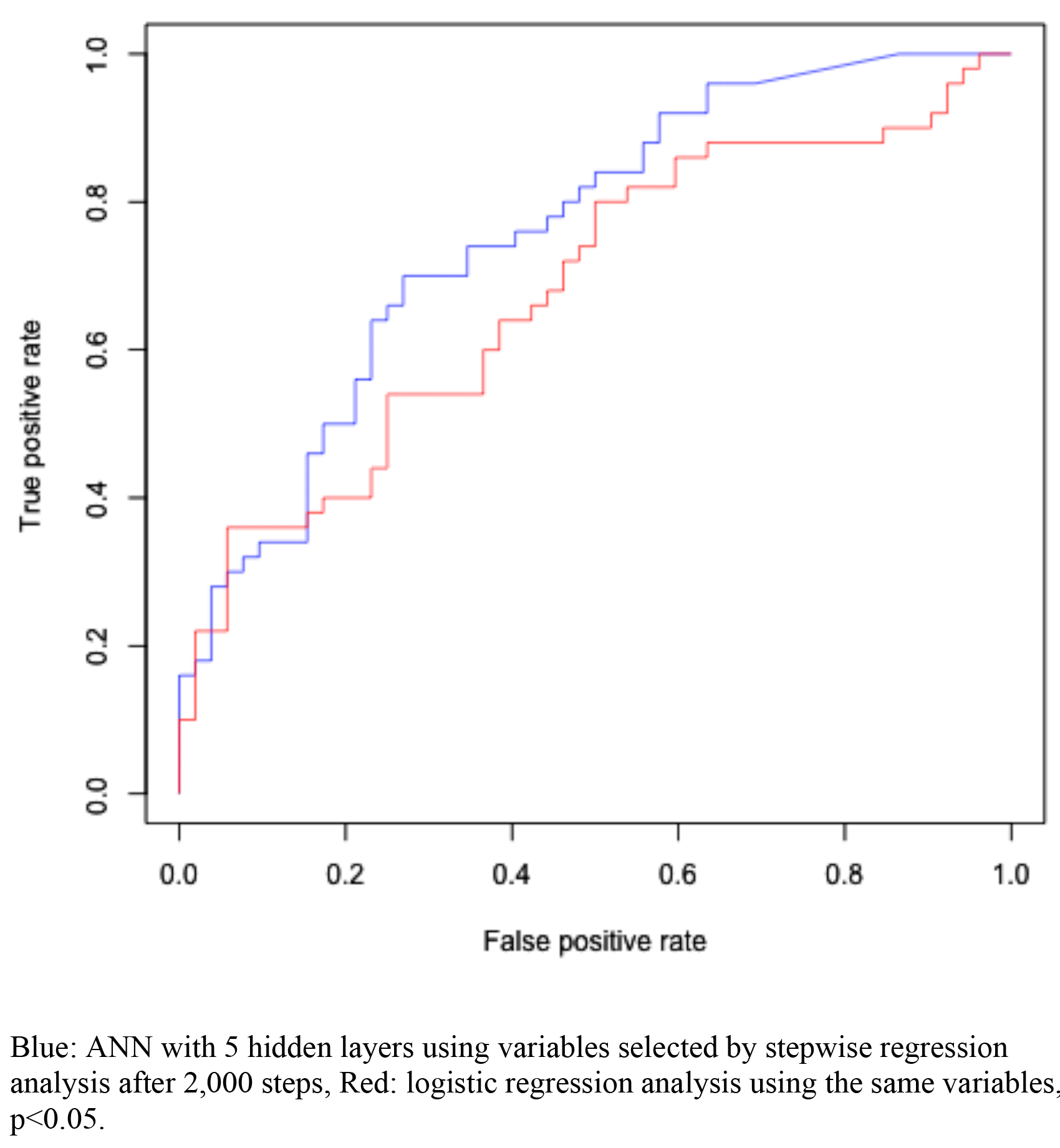
ROC curves for the actual detection of prostate cancer on biopsy to assess the usefulness of the algorithm-output probability of prostate cancer in test samples.

**Fig. 3:**
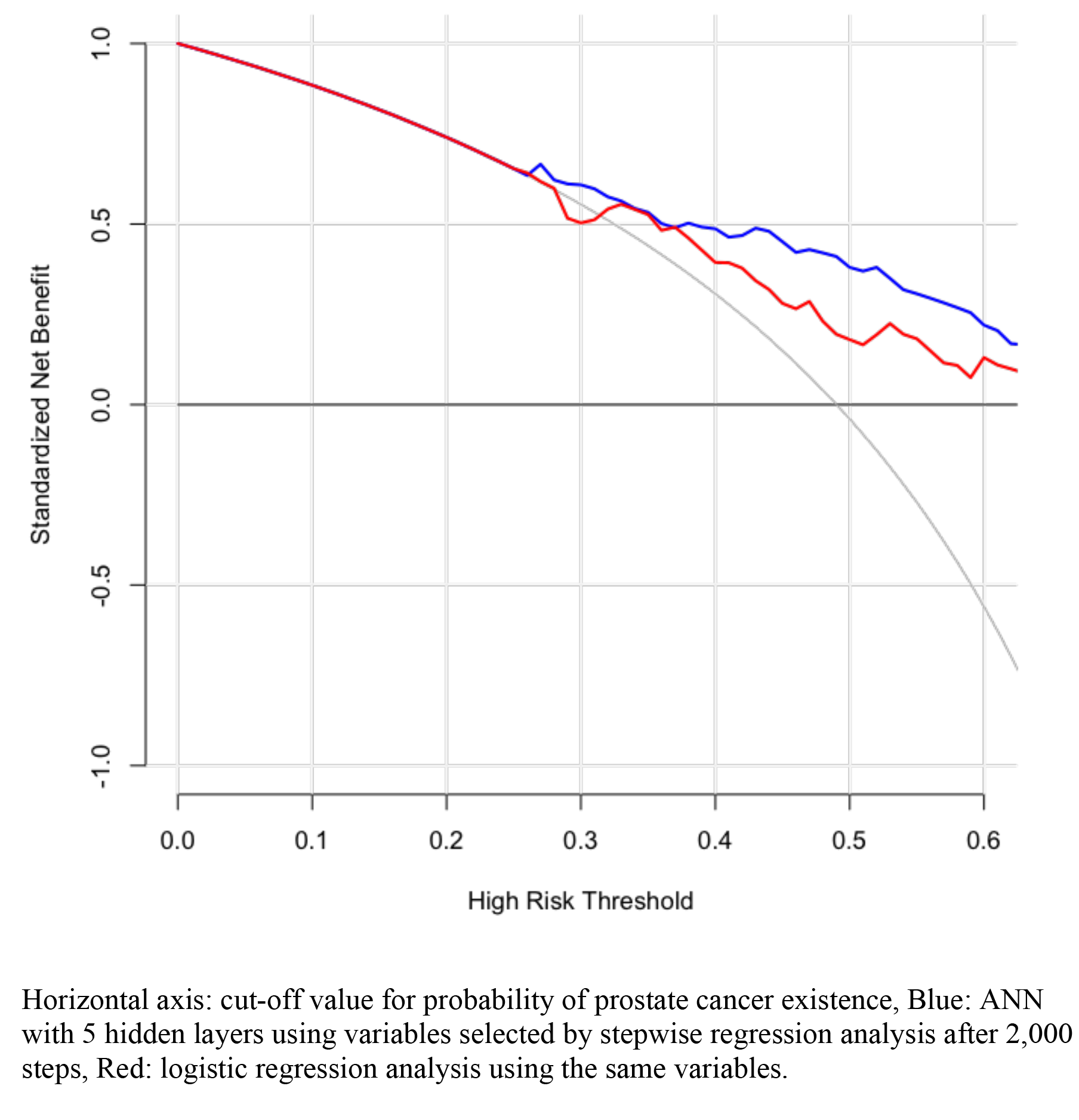
Net-benefit curves for any prostate cancer detection.

**Table 5:**
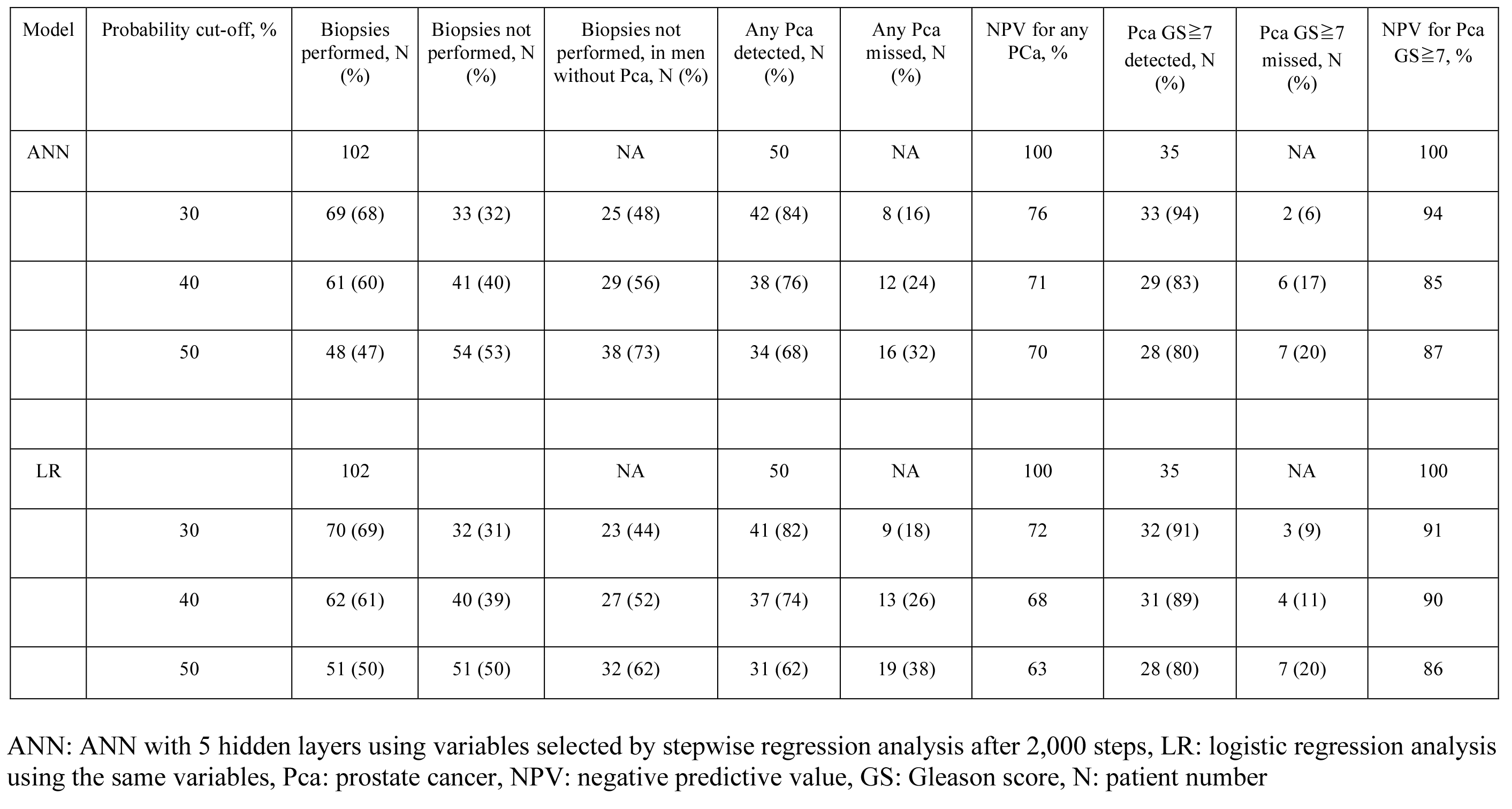
Comparison of performance of ANN and the corresponding LR.

## Discussion

Multilayer ANN could predict prostate cancer with a 5-10% higher accuracy than conventional LR when used as a discrete classifier at a cut-off probability of 0.5. This superiority was also supported by the fact that the AUCs with ANN, of which the parameter was the probability of prostate cancer existence, were larger than with LR when used as a probabilistic classifier. Thus, multilayer ANN may be a promising tool to predict the existence of prostate cancer.

However, with precise investigation, the difference between ANN and LR was still marginal for the prevention of unnecessary prostate biopsy, missing prostate cancer, and negative predictive value. Accurate prediction of prostate cancer with a high Gleason score (GS) is clinically more important, as it tends to become castration-resistant, metastasize, and become life-threatening. The marginal difference also applies to high-GS (≥7) prostate cancer.

Trials to predict the presence of prostate cancer without prostate biopsy have been conducted using LR by many investigators (8–15), leading to excellent performance. LR is already very accurate in its prediction of prostate cancer and it may retain this effectiveness even on comparison with ANN. ANN and LR missed less than 20% of all prostate cancer and less than 10% of high-GS prostate cancer on setting the cut-off value of probability as 0.3. This rate of missed cancer detection may be acceptable or remain controversial when ANN and LR are used to screen candidates for prostate biopsy.

The ANN predicting the existence of prostate cancer is novel and still preliminary. The accuracy of predicting the existence of prostate cancer without biopsy is good, but there is still room for improvement. The highly effective ANN shows a higher rate of classification accuracy and larger AUC. There are ways to improve the capacity of ANNs to predict prostate cancer.

Increasing the sample size may improve the performance of multilayer ANN as a result of more sufficient training. In this study, the sample size was limited and training with 10 to 100 times more samples would work better. If the sample size is increased, ANNs with more hidden layers and neural nodes can perform better, avoiding early over-fitting. Adding more variables that are appropriate for predicting prostate cancer may be another good way for ANN to perform better. For example, genetic factors and excellent biomarkers may be candidates in addition to standard clinical and imaging parameters. Tuning of hyper-parameters such as the learning rate, regularization penalty scale, and leaning cycles would be less effective than those factors described above. In conclusion, ANN accurately predicted prostate cancer without biopsy marginally better than LR. However, Still, for clinical application, ANN performance may still need improvement.

